# Host body size, not host population size, predicts genome-wide effective population size of parasites

**DOI:** 10.1101/2022.10.06.511102

**Authors:** Jorge Doña, Kevin P. Johnson

**Author notes:** Jorge Doña and Kevin P. Johnson., **Email:** Jorge Doña; Kevin P. Johnson. **Competing Interest Statement:** None.

## Abstract

The effective population size (*N*_*e*_) of an organism is expected to be generally proportional to the total number of individuals in a population. In parasites, we might expect the effective population size to be proportional to host population size and host body size, because both are expected to increase the number of parasite individuals. However, among other factors, parasite populations are sometimes so extremely subdivided that high levels of inbreeding may distort these predicted relationships. Here, we used whole-genome sequence data from dove parasites (71 feather louse species of the genus *Columbicola*) and phylogenetic comparative methods to study the relationship between parasite effective population size and host population size and body size.We found that parasite effective population size is largely explained by host body size but not host population size. These results suggest the potential local population size (infrapopulation or deme size) is more predictive of the long-term effective population size of parasites than is the total number of potential parasite infrapopulations (i.e., host individuals).

**Impact Summary:** Parasites, among Earth’s most diverse, threatened, and under-protected animals, play a central role in ecosystem function. The effective population size (*N*_*e*_) of an organism has a profound impact on evolutionary processes, such as the relative contributions of selection and genetic drift to genomic change. Population size is also one of the most important parameters in conservation biology. For free-living organisms, it is expected that *N*_*e*_ is generally proportional to the total number of individuals in a population. However, for parasites, among other factors, populations are sometimes so extremely subdivided that high levels of inbreeding may distort these relationships. In this study, we used whole-genome sequence data from dove parasites and phylogenetic comparative methods to investigate the relationship between parasite effective population size (*N*_*e*_) and host population size and body size. Our results revealed a positive relationship between parasite effective population size (*N*_*e*_) and host body size, but not host population size. These results suggest that the size of parasite infrapopulations may be the most important factor when considering parasite effective population size, and have important implications for conservation.

## Introduction

The effective population size (*N*_*e*_) of an organism has a profound impact on evolutionary processes, such as the relative contributions of selection and genetic drift to genomic change (Wright, 1943; Waples, 2002; Charlesworth, 2009). For free-living organisms, it is generally expected that *N*_*e*_ is proportional to the total number of individuals in a population (census population size, *N*_*c*_) (Frankham, 1995; Waples, 2002). While population size estimates can often be readily obtained for free-living species, estimating the population size of parasites can be more challenging because this usually requires sampling from individual hosts (Criscione and Blouin 2005, Criscione et al., 2005; Poulin, 2007; Clayton et al., 2015; Strobel et al., 2019).

The relationship between long-term effective population size (*N*_*e*_) and the total number of individuals in a population (*N*_*c*_) is complex and depends on various ecological and demographic factors (Buffalo, 2021; Charlesworth & Jensen, 2022). According to neutral theory, genetic diversity is expected to increase with population size (Kimura, 1971). However, the observed levels of diversity across metazoans vary only two orders of magnitude while population sizes vary over several orders of magnitude, known as Lewontin’s Paradox of Variation (1974). The reasons for this discrepancy remain unclear (Buffalo, 2021; Charlesworth & Jensen, 2022), making it challenging to establish a strong relationship between *N*_*e*_ and *N*_*c*_ using population size estimates alone.

Considering the complex interplay of factors that influence *N*_*e*_, an optimal approach to studying the effect of any given factor on *N*_*e*_ involves holding other factors constant as much as possible. Ecological replicate systems, defined as species sharing similar natural histories, are particularly valuable for investigating the relationship between complex variables (Clayton et al., 2015). Host-parasite systems provide an opportunity to hold many other factors constant when making comparisons across parasite species. Such ecological replicate systems have facilitated research on identifying factors influencing the evolutionary history of parasites, e.g., the influence of dispersal capabilities on coevolutionary history (Clayton & Johnson, 2003; Harbison & Clayton, 2011; Doña et al., 2020). Thus, while it may not be possible to quantify all possible parameters that affect *N*_*e*_ in these systems, it is reasonable to assume that they are generally similar across species in such systems. This assumption is based on the fact that ecologically similar parasites have comparable life cycles and modes of transmission and inhabit very similar host species.

Typically, we might expect that the size of a parasite population is proportional to that of the host, because parasites rely on their hosts for survival and reproduction (Poulin, 2007; Barrett et al., 2008; Clayton et al., 2015). However, population subdivision can also influence measures of *N*_*e*_ for a species (Wright, 1943; Charlesworth et al., 2003). For example, theoretical expectations, based on the island model of population structure, generally predict that subdivided populations have a higher overall long-term *N*_*e*_ than non-subdivided ones (Charlesworth et al., 1997; Charlesworth et al., 2003; Charlesworth, 2009). On the other hand, in highly divided populations, levels of inbreeding can increase, such that the local *N*_*e*_ of these individual subdivided populations can be low (Charlesworth et al., 1997; Charlesworth et al., 2003; Charlesworth, 2009). Here, we are concerned with measures of long-term *N*_*e*_ as predicted by coalescent theory (*θ=*4*N*_*e*_*μ*).

In the case of parasites, the concept of ‘population’ differs from that of free-living organisms. Populations of parasites that live on a host can often be readily demarcated. Parasite populations can sometimes be so extremely subdivided that each individual host harbors a distinct infrapopulation (groups of parasites of the same species living in a single host), analogous to a deme (Bush et al., 1997; Huyse et al., 2005; Criscione and Blouin 2005, Criscione et al., 2005; Poulin, 2007; Clayton et al., 2015). This subdivision is particularly pronounced in the case of parasites that spend their entire life-cycle on the host (i.e., permanent parasites, DiBlasi et al., 2018; Sweet and Johnson, 2018; Virrueta-Herrera et al., 2022). Lice, which are permanent parasitic insects of birds and mammals, have highly structured infrapopulations subject to high levels of inbreeding (Virrueta-Herrera et al., 2022). Because it can generally be assumed that the census size of a parasite species is reflected by the mean infrapopulation size multiplied by the number of infrapopulations (Criscione et al., 2005; Doña et al., 2015), we might expect infrapopulation size to be correlated with the effective population size of a parasite, as has been shown for feather mites (Doña et al., 2015).

Host body size has been shown to strongly impact infrapopulation size, with larger-bodied hosts harboring larger parasite infrapopulations (Poulin, 1999; Poulin, 2007; Clayton et al., 2015). For example, a positive effect of host body size on parasite abundance has been shown for avian feather lice, which feed on the feathers of their hosts (Rozsa, 1997; Clayton and Walther, 2001). Accordingly, we would expect that feather lice on larger-bodied avian hosts would have higher *N*_*e*_, reflecting their larger infrapopulation sizes, and perhaps higher overall population size.

Thus, host population size and host body size are two factors that may influence parasite’s *N*_*e*_. Here, we test the relative contributions of these two factors to parasite *N*_*e*_ by examining genome-wide variation in the wing lice (Phthiraptera: *Columbicola*) of pigeons and doves (Aves: Columbidae). Pigeons and doves vary dramatically in overall population sizes, with some species being among the most abundant birds on earth and others restricted to single small islands and highly endangered. In addition, pigeons and doves vary by over an order of magnitude in body mass, and smaller-bodied species have been shown to have smaller infrapopulations of these lice (Rozsa, 1997). Thus, pigeons and doves and their lice are an excellent system in which to examine the correlation between both host population size and host body size and parasite *N*_*e*_, while holding many other ecological variables constant. We used genome sequencing of 71 species of *Columbicola* to estimate a phylogeny for these parasites and examine the relationship between a genome-wide measure of effective population size (*θ=*4*N*_*e*_*μ*) and the overall population size and body size of their respective hosts, accounting for parasite phylogeny.

## Materials and Methods

### Taxon sampling and host data

We sampled 94 individual lice, representing 71 different species of *Columbicola* (Table S1), which are feather lice (Insecta: Ischnocera) of pigeons and doves. We also included five feather louse outgroup taxa for the phylogenomic analyses, selected based on recent higher level phylogenomic studies of feather lice (Table S1). We obtained host body size (body mass) information from the Birds of the World online database (Billerman et al., 2022). Specifically, in cases where measures from both males and females were reported independently, we used the average between male and female body mass. We obtained global-scale host population size data from recent estimates (Callaghan et al., 2021), which estimated species-specific abundances for approximately 92% of all extant bird species by integrating data from well-studied species with a global dataset of bird occurrences throughout the world. The authors used missing data theory to estimate species-specific abundances and their associated uncertainties. In particular, we used the “Abundance estimate” data from the “Dataset_S01.xlsx” supplemental file.

### Genomic sequencing

Some of the genomic data we analyzed here have been previously published (Boyd et al., 2017, see Table S1 for details). For the newly sequenced samples, which had been stored in 95% ethanol at −80 °C, we performed single-louse DNA extractions and photographed each specimen as a voucher. We extracted total genomic DNA by first letting the ethanol evaporate and then grinding the louse with a plastic pestle in a 1.5 ml tube. For DNA extraction, we used a Qiagen QIAamp DNA Micro Kit (Qiagen, Valencia, CA, USA) and conducted an initial incubation at 55 °C in buffer ATL with proteinase K for 48 h. Otherwise, we followed the manufacturer’s protocols and eluted purified DNA off the filter in a final volume of 50ul buffer AE. We used a Qubit 2.0 Fluorometer (Invitrogen, Carlsbad, CA, USA) and the high sensitivity kit to quantify total DNA.

We prepared genomic libraries using the Hyper library construction kit (Kapa Biosystems). We then sequenced these libraries to generate 150 bp paired-end reads using Illumina NovaSeq 6000 with S4 reagents. Libraries were tagged with unique dual-end adaptors and multiplexed 48 libraries per lane, intending to achieve approximately 30-60X coverage of the nuclear genome. We trimmed adapters and demultiplexed the sequencing data using bcl2fastq v.2.20 to generate final fastq files. We deposited raw reads for each library in NCBI SRA (Table S1).

### Single-copy orthologs assembly

We used fastp v0.20.1 (Chen et al., 2018) to perform adaptor and quality trimming (phred quality >= 30). We then converted trimmed fastq files to aTRAM 2.0 blast databases using the atram_preprocessor.py command of aTRAM v2.3.4 (Allen et al., 2018). We used an amino acid sequence reference set of 2395 single-copy ortholog protein-coding genes (Johnson et al., 2018) from the human louse, *Pediculus humanus* (Kirkness et al., 2010). We assembled the single-copy ortholog genes using the atram.py command and the ABySS assembler with the following parameters (iterations = 3, max-target-seqs = 3000). The Exonerate pipeline in aTRAM (atram_stitcher.py command) was used to stitch together exon sequences from these protein-coding genes (Slater and Birney, 2005).

### Species delimitation

Several prior studies have indicated the potential for cryptic species within species of *Columbicola* (Johnson et al., 2007; Malenke et al., 2009; Sweet and Johnson, 2018), and we wanted to account for this in our comparative analyses. For assembly of the mitochondrial COI gene, we subsampled four million reads (two million read1 and two million read2) from each library using Seqtk v1.3 (Li, 2022). As the reference target for constructing COI sequences from all samples in our current work, we used a COI sequence from *Columbicola columbae* that had previously been published (Johnson et al., 2007). For these assemblies, we ran aTRAM for only a single iteration. Then, we translated COI DNA sequences to amino acids, aligned them, and threaded the DNA back through the aligned proteins to obtain nucleotide data. As a quality control procedure, we blasted COI sequences against NCBI to identify any identical or nearly identical to previously generated Sanger sequences. We estimated a phylogenetic tree based on these COI sequences under maximum likelihood using model parameters estimated by IQ-TREE 2 v.2.1.235 (Minh et al., 2020). The best model chosen for our analysis was K3Pu+F+I+G4, selected based on the Bayesian Information Criterion (BIC). The optimal proportion of invariable sites (pinv) and gamma shape parameter (alpha) were 0.277 and 0.490, respectively, with a LogL of -51235.155. We estimated ultrafast bootstrap support values with UFBoot2 (Hoang et al., 2017). Finally, we also computed the percent pairwise sequence divergences among all the COI sequences (using the R function dist.dna, model “raw,” pairwise.deletion = T from APE v5.5, Paradis and Schliep, 2018) and looked at their distribution to identify likely cryptic species, which indicated a 5% uncorrected p-distance threshold would be appropriate, as in prior studies of lice (Johnson et al., 2021).

### Generation of a dated ultrametric phylogenetic tree

#### Phylogenomic analyses

We translated assembled single-copy-ortholog nucleotide sequences to amino acids and aligned them using MAFFT v.7.47133 (Katoh and Standley, 2013). After threading the DNA back through the aligned proteins to obtain nucleotide alignments, we used trimAL v.1.4.rev2234 (with a 40% gap threshold) (Capella-Gutiérrez et al., 2009) to trim individual gene alignments. We discarded any gene present in less than four taxa. We then concatenated gene alignments into a supermatrix and analyzed it under maximum likelihood using IQ-TREE 2 in a partitioned analysis that included model selection for each partition. Support was estimated using ultrafast bootstrapping (Hoang et al., 2017). We also ran a coalescent analysis using ASTRAL-III (Zhang et al., 2018) on individual gene trees estimated by maximum likelihood in IQ-TREE 2. As a measure of branch support, we computed local posterior probability for each branch in ASTRAL-III. Both trees were almost identical; therefore, we only used the partitioned concatenated tree for dating and phylogenetic comparative analyses.

#### Obtaining calibration points from cophylogenetics

We used eMPRess v1.0 (Santichaivekin et al., 2020) to compare host and parasite trees based on their topology. As in prior cophylogenetic studies, we used costs of duplication: 1, sorting: 1, and host-switching: 2. This is the cost scheme used by most published cophylogenetic studies of lice, as well as other groups of ectosymbionts (Doña et al., 2017; Matthews et al., 2018; Sweet and Johnson, 2018; Johnson et al., 2021, 2022; Boyd et al., 2022). For the host tree, we obtained phylogenetic information from a prior phylogenomic study (Boyd et al., 2022). As there was no phylogenetic information for fourteen of the focal species in this tree, we obtained the placement of these species from additional phylogenetic studies (Johnson and Weckstein, 2011; Sweet et al., 2017; Nowak et al., 2019), and used the R function bind.tip from phytools v.1.2-0 (Revell, 2012) to incorporate these species into our tree (See Figure S1). We used the phylogeny derived from the partitioned analysis (above) for the parasite tree. Based on the distribution of the MPR distances histogram, we summarized the MPR space into one cluster and drew a representative median MPR. From this reconstruction we identified terminal cospeciation events between sister pairs of doves and lice to use in the molecular dating analysis (below).

#### Dating analysis

We produced an ultrametric tree using the least square dating (LSD2) method implemented in IQ-TREE (To et al., 2016). Because there are no currently known fossilized lice within Ischnocera, we used terminal cospeciation events between sister pairs of doves and lice (above) as calibration points for molecular dating (Johnson et al., 2021, 2022). Specific cospeciation events that were used as calibration points can be found at Table S2 (see Supplemental information). For this analysis, we set a root age of 52 mya (based on de Moya et al., 2019) as required by the LSD2 software, which needs at least one fixed calibration point to obtain a unique optimal solution for the time estimates. We also set a minimum branch length constraint (u = 0.01) to avoid collapsing short but informative branches without introducing bias to the time estimates (see https://github.com/tothuhien/lsd2). The resulting ultrametric tree (Figure S2) was used for phylogenetic generalized least squares (PGLS) analyses to investigate the correlation between the dependent parasite variable *θ* (a measure of parasite *N*_*e*_) and the independent variables of host population size and host body size.

### SNP calling and mlRho analyses

We used the *Columbicola columbae* chromosome-level genome assembly (Baldwin-Brown et al., 2021) as the reference for the SNP calling analyses. We aligned trimmed and filtered reads to the *C. columbae* reference genome using bwa v0.7.17 (Li and Durbin, 2009). We then removed PCR duplicates with picard v2.26.10 (Broad Institute, 2022) and sorted and indexed bam files with samtools v1.14 (Danecek et al., 2021). We called SNPs using bcftools multiallelic caller (Danecek and McCarthy, 2017). Lastly, we used vcftools to filter the vcf file with the following filtering parameters: <40% missing data, site Phred quality score >30, a minimum genotype depth of 10X, a maximum genotype depth of 60X, a minimum mean site depth of 10X and a maximum mean site depth of 60X. A total of 177,895 SNPs remained after filtering.

We used mlRho v2.9 (Haubold et al., 2010) to calculate the sample-specific mean theta (*θ*), which is defined as the population mutation rate, or *θ* = 4*N*_*e*_*μ*, and which can be used as an indicator of long-term effective population size (Lynch, 2008; Meyer et al., 2012; Virrueta-Herrera et al., 2022) because it is proportional to *N*_*e*_. This approach, based on coalescent theory, provides an average measure of the effective population size over a relatively long period of time, incorporating the influence of historical fluctuations in population size. This method also helps address potential biases arising from unequal sequence coverage, where there is a relatively high probability that all sequences at a locus will be derived from just one of the two parental chromosomes in a diploid individual, or from sequence errors. The method assumes a constant mutation rate (*μ*) and a constant population size over time, although it can accommodate minor fluctuations in these parameters. Thus, our estimates of long-term effective population size should provide a robust and informative measure for examining the relationships between parasite *N*_*e*_, host population size, and host body size in a comparative framework of ecological replicates. For this analysis, we converted bam files from bwa to profile (.pro) files for each individual louse and then ran mlRho with maximum distance (M) = 0.

### Phylogenetic comparative methods

We used phylogenetic generalized least squares (PGLS) models, gls function from nlme v3.1-149 R package (Pinheiro et al., 2020), to examine associations between *θ* (a measure of parasite *N*_*e*_) and host population size and host body size. We used the obtained dated ultrametric tree as the phylogenetic tree for all PGLS analyses, evaluated various phylogenetic correlation structures in our regressions (corPagel, corBrownian), and used AIC model comparisons to identify the best fitting correlation structure for the models. We fitted PGLS models with the following formulas: Parasite_θ ∼ HostBodySize, Parasite_θ ∼ HostPopulationSize, and Parasite_θ ∼ HostBodySize + HostPopulationSize. For each of these formulas, we tried both corPagel and corBrownian correlation structures, resulting in a total of six models. CorPagel accounts for a variable rate of phylogenetic signal (Pagel, 1999; Freckleton et al., 2002), whereas corBrownian assumes a constant rate of trait evolution along the branches of the phylogenetic tree (Felsenstein, 1985; Martins & Hansen, 1997). We checked models via visual inspection of diagnostic plots (residuals vs. fitted values and QQ plots to check normality).

## Results

Out of the initial 94 samples, 18 were excluded from further analyses because they had mitochondrial COI genetic distances less than 5% in the species delimitation analysis. The same result was on inspection of the COI tree, in which these 18 samples were grouped with the retained species with zero or almost zero branch length, indicating that they likely represent the same species. The topologies of both phylogenomic trees based on the 2395 target single copy nuclear gene orthologs were nearly identical, with high support in the concatenated partitioned analysis (86 out of 92 nodes had 100% support values, 93%) and the coalescent tree (87 out of 93 nodes had posterior probabilities equal to 1, 94%). In the cophylogenetic analysis, we found that the reconstruction had a lower cost value that expected by chance (p-value=0.009901), indicating significant congruence likely owing to cospeciation between hosts and parasites.

Specifically, we identified 36 cospeciation events, 7 of which occurred between terminal sister species and were used to generate the dated ultrametric tree for the PGLS analysis.

In comparative analysis of effective population size across species of *Columbicola*, we found a strong positive relationship between *θ*, a metric directly proportional to *N*_*e*_, and host body size (PGLS, Brownian: R^2^_pred_ = 0.44, p = 0.0005; Pagel’s λ: R^2^_pred_ = 0.48, p = 0.0004; Fig. 1). In contrast, there was no significant relationship between *θ* and host population size (PGLS, Brownian, p = 0.0908; Pagel’s λ, p = 0.09). Including host population size in the best model led to a small improvement in the overall model fit (PGLS, Pagel’s λ including host population size, R^2^_pred_ = 0.52), but the host population size term remained non-significant (PGLS, Pagel’s λ model including host population size, p = 0.1454). We further confirmed the lack of relationship between host population size and *θ* by performing a simple linear regression analysis (not accounting for parasite phylogeny), which showed no significant relationship between the two variables (R^2^ = 0.0013, p = 0.92; see Figure S3 in the Supplementary Material). These results are consistent with the finding above, which took into account phylogeny, that no significant relationship exists between *θ* and host population size across the entire dataset.

**Figure 1.**
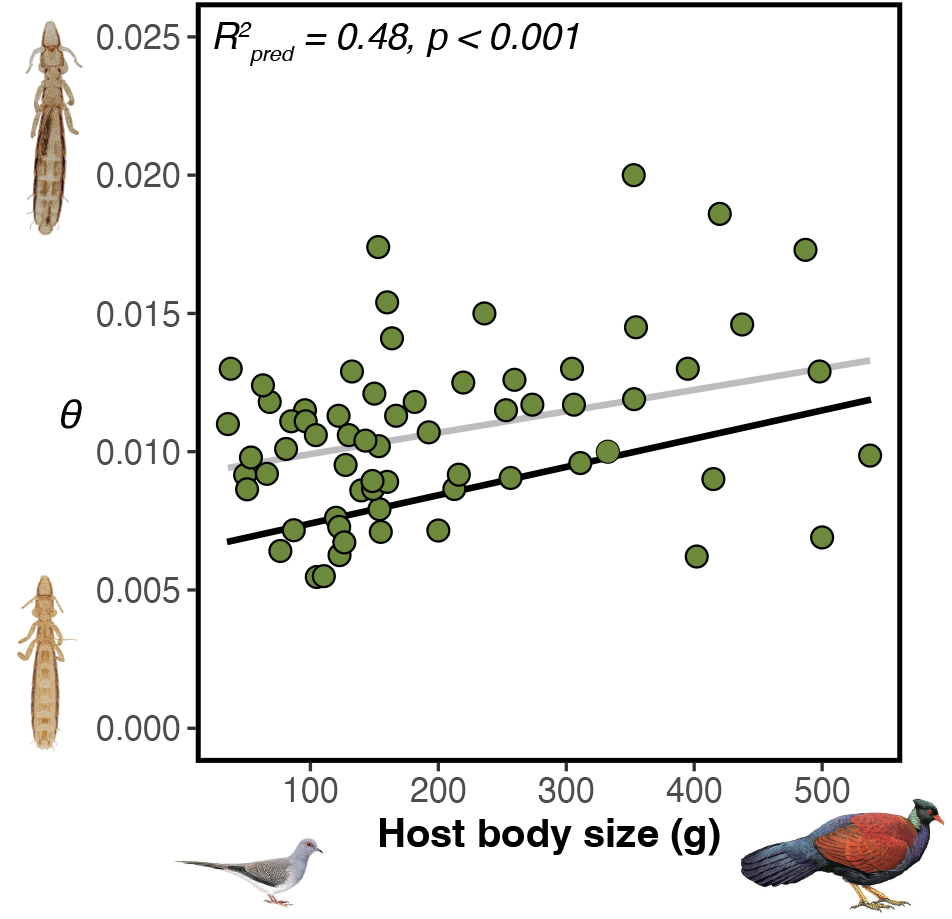
Relationship between a genome-wide measure (*θ*) of effective population size and host body size. The black regression line corresponds to the PGLS model and the gray regression line to the same GLS model without accounting for phylogenetic non-independence. Credit: louse photos on the left: Stephany Virrueta-Herrera; bird illustrations on the bottom, ©Lynx Edicions (*Otidiphaps nobilis*: Hilary Burn; *Geopelia cuneata*: Martin Elliott).

## Discussion

For parasites such as lice, hosts represent their habitat (Clayton et al., 2015). Host body size largely explains parasite infrapopulation (deme) size (Rozsa, 1997; Clayton and Walther, 2001). Genome scale data for parasitic lice of pigeons and doves revealed that metrics (*θ*), associated with effective long-term population size (*N*_*e*_), are also highly correlated with host body size. In contrast, there was little association between parasite effective population size and host population size. Thus, it appears that the larger infrapopulation sizes of parasites on larger-bodied hosts may directly influence long-term *N*_*e*_ by a direct correlation between parasite infrapopulation size and overall parasite population size, eliminating any effects of overall parasite population size as related to host population size. However, prior studies have also indicated that the relationship between census population size and long-term effective population size may not typically be strong (Buffalo, 2021; Charlesworth & Jensen, 2022). Thus, our finding of no relationship between effective population size of parasites and census population size of their hosts may not be entirely unexpected. In addition, other factors may also influence our estimator of *N*_*e*_, besides overall population size alone. Despite these considerations regarding interpretation, the general pattern in our results was quite clear. The comparative nature of our study, which includes multiple ecological replicates, and the consistent application of data and bioinformatic analyses across species serve as strengths in assessing the relationship between host body size and parasite *N*_*e*_.

Despite the possibility that infrapopulation size is directly correlated with overall population size of parasites, there are also other factors to consider in these host-parasite systems. Several studies have indicated that louse infrapopulations on single host individuals are highly inbred, showing strong evidence of genetic structure even between host individuals in close proximity (Ascunce et al., 2013; DiBlasi et al., 2018; Virrueta-Herrera et al., 2022). This inbreeding would reduce the effective population size within demes on single host individuals. However, theoretical models predict that population structure should increase the long-term effective population size (which is what we measured) based on coalescence (Charlesworth et al., 1997; Charlesworth et al., 2003; Charlesworth, 2009), because alternative alleles can go to fixation in different infrapopulations increasing the overall standing genetic diversity of the global population. Counter to this expectation, we find that the estimator of *N*_*e*_ is lower for parasites on small-bodied doves that are expected to host smaller local infrapopulations with higher levels of inbreeding, which would result in more structure among demes.

Another factor to consider is that smaller-bodied host species also typically have a lower parasite prevalence (i.e., proportion of host individuals that are inhabited by the parasite) (Bush et al., 1997). This pattern might be due to smaller infrapopulations being more susceptible to local extinction because of environmental and demographic stochasticity, a known factor shaping *N*_*e*_ (Charlesworth, 2009; Doña and Johnson, 2020). Therefore, host body size could influence local extinction probability of parasites and thus play a role in determining long-term *N*_*e*_ of permanent parasites (Farrell et al., 2021). Given the lower prevalence and intensity of lice on small-bodied hosts, it may be that the total number of lice in the global population is considerably smaller than those found on large-bodied hosts. While it might be expected that small-bodied doves have generally larger population sizes, because of the general inverse relationship between body size and population size of most organisms (White et al., 2007), we found no such relationship in our dataset (R^2^ = 0.003, p > 0.1). This finding agrees with previous results on other birds (Nee et al., 1991). Thus, while further research on global population estimates of louse species would help understand these relationships, our results suggest that at lower taxonomic levels, host body size and not host population size is the most explanatory factor of parasite *N*_*e*_. However, one recent study has shown that even a moderate shift in host specificity (which is expected to correlate with overall host population size) can translate into significant differences in genetic characteristics of parasite populations (Martinů et al., 2018), so this topic deserves further study.

Another factor to consider is the low among-deme migration rates of permanent parasites. A low migration rate among parasite infrapopulations is expected to increase long-term *N*_*e*_ because of increased population substructuring. Permanent parasites, such as lice, have minimal dispersal capabilities and thus migration rates are expected to be very low. While host population size has been previously identified as a potential driver of parasite population dynamics (Doña and Johnson, 2020), the lack of relationship between parasite *N*_*e*_ and host population size might be indicative of these very low migration rates. In this case, a single migrant contributes more to homogenizing infrapopulations on small-bodied hosts than it does to reducing infrapopulation structure of lice on large bodied hosts, because of differences in the probability of fixation of migrant alleles in large and small populations. This factor could contribute to our observation of higher *N*_*e*_, perhaps owing to more population structure, of parasite populations on large-bodied hosts.

In our analysis, we also did not account for the potential effects of selection or demography separately, and assumed that these factors are largely similar across species, allowing us to make meaningful comparisons. However, selection is also known to influence effective population size (Charlesworth et al., 1997; Charlesworth et al., 2003; Charlesworth, 2009). For loci under selection, the realized effective population size is lower than those whose frequency is only affected by drift (Charlesworth et al., 1997; Charlesworth et al., 2003; Charlesworth, 2009).

Louse species with smaller infrapopulation sizes, and higher inbreeding, might suffer more from inbreeding depression. This would be a genome wide negative selection, which would be predicted to lower overall effective population size (Hedrick and García-Dorado, 2016). However, the power of genome wide negative selection to eliminate variant deleterious alleles would also be stronger in larger populations, such as those on large-bodied hosts, and thus the predicted correlation between *N*_*e*_ and host body size would be opposite to what we observed. It is unclear how these two opposing forces, inbreeding depression versus the elimination of deleterious alleles by selection, might play out under this situation. In addition, some studies suggest that species under similar conditions to lice (small populations, fluctuating in size) usually do not suffer from inbreeding depression, due to the purging of deleterious alleles (Larsen et al., 2011; Robinson et al., 2018; Valk et al., 2021). It is unknown if lice suffer from inbreeding depression, given that they normally experience high levels of inbreeding, but would be a topic of interest for future investigation.

Considerations of effective population size also have implications for conservation. Parasites are among the earth’s most diverse, threatened, and under-protected animals (Carlson et al., 2017). Under the global parasite conservation plan, risk assessment, along with applying conservation genomics to parasites, were identified as two of the major goals for parasite conservation over the next decade (Carlson et al., 2020). Our result that host body size, but not host population size, is a good predictor of parasite *N*_*e*_ can easily translate into parasite conservation practices, drawing attention to conservation of smaller bodied hosts as a practice to conserve parasites.

Overall, our study shows that host body size plays a major role in shaping parasite population genomics and provides evidence for the essential role that individual hosts play as habitat for permanent parasites with very limited transmission abilities. Despite potential complexities arising from factors such as inbreeding and colonization dynamics, the comparative nature of our study, which includes multiple ecological replicates, ensures some level of robustness of our overall conclusions. As we continue to investigate host-parasite interactions, understanding the role of host traits in shaping parasite population genomics will become increasingly important, especially for developing informed conservation strategies for these diverse and threatened organisms

## Acknowledgments

We thank J.M. Bates, B. Benz, S.E. Bush, D.H. Clayton, T. Chesser, R. Faucett, R. Moyle, V.Q. Piacentini, F. Sheldon, A.D. Sweet, J.D. Weckstein, and B. Zonfrillo for assistance in obtaining specimens. We thank B.M. Boyd and S. Virrueta Herrera for assistance with gDNA extractions. A. Hernandez and C. Wright at the University of Illinois Roy J. Carver Biotechnology Center provided assistance with Illumina sequencing. We thank K.K.O. Walden for assistance with submitting reads to NCBI SRA. Funding was provided by US NSF DEB-1342604, DEB-1925487 and DEB-1926919 grant awards to K.P.J., and European Commission grant H2020-MSCA-IF-2019 (INTROSYM:886532) to J.D. We also thank reviewers for their constructive comments.

## Author Contributions

J.D. designed the study, conducted the analyses, prepared figures, wrote the manuscript draft, and edited the manuscript. K.P.J. designed the study, obtained funding, wrote the manuscript draft, and edited the manuscript.

## Data accessibility

Intermediary files generated in this study have been deposited in Figshare (reserved DOI: 10.6084/m9.figshare.21269640; private link for review: https://figshare.com/s/2f2de5dc909155da815a).

## Notes

https://figshare.com/s/2f2de5dc909155da815a

### Competing Interest Statement

The authors have declared no competing interest.

### Summary of Updates

In this revised version of the manuscript, we have addressed the reviewers' comments and made several important updates to enhance the clarity and robustness of our study. The main changes include: 1. We expanded the explanation of our methods for estimating effective population size (Ne) and discussed the implications of our approach. 2. We clarified our use of the Columbicola columbae genome as a reference for mapping reads and provided justification for this choice. 3. We added information on the mlRho method for estimating theta and acknowledged potential limitations in the Discussion. 4. We provided a clearer description of the PGLS models fitted in our study and discussed the potential non-independence of observations of parasite diversity due to host phylogeny. 5. In response to minor comments, we made several adjustments to the text, such as providing more information on data sources, clarifying our methodology for obtaining nucleotide alignments, and reporting actual p-values for our PGLS analysis. 6. We have included the dated ultrametric louse phylogenies and the bird tree in the supplementary material, as well as uploaded the host tree to the study's Figshare repository.

## References

Allen, J.M., LaFrance, R., Folk, R.A., Johnson, K.P. & Guralnick, R.P. (2018) aTRAM 2.0: An Improved, Flexible Locus Assembler for NGS Data. Evolutionary Bioinformatics, 14, 1176934318774546.

Ascunce, M.S., Toups, M.A., Kassu, G., Fane, J., Scholl, K. & Reed, D.L. (2013) Nuclear Genetic Diversity in Human Lice (Pediculus humanus) Reveals Continental Differences and High Inbreeding among Worldwide Populations. PLoS ONE, 8, e57619.

Baldwin-Brown, J.G., Villa, S.M., Vickrey, A.I., Johnson, K.P., Bush, S.E., Clayton, D.H., et al. (2021) The assembled and annotated genome of the pigeon louse Columbicola columbae, a model ectoparasite. G3 Genes|Genomes|Genetics.

Barrett, L.G., Thrall, P.H., Burdon, J.J. & Linde, C.C. (2008) Life history determines genetic structure and evolutionary potential of host–parasite interactions. Trends in Ecology & Evolution, 23, 678–685.

Billerman S. M., Keeney, B. K., Rodewald, P. G., and Schulenberg T. S. (2022) Birds of the World. Cornell Laboratory of Ornithology, Ithaca, NY, USA.

Boyd, B.M., Allen, J.M., Nguyen, N.-P., Sweet, A.D., Warnow, T., Shapiro, M.D., Villa, S.M., Bush, S.E., Clayton, D.H. & Johnson, K.P. (2017) Phylogenomics using target-restricted assembly resolves intra-generic relationships of parasitic lice (Phthiraptera: Columbicola). Systematic Biology, 66, 896–911.

Boyd, B.M., Nguyen, N.-P., Allen, J.M., Waterhouse, R.M., Vo, K.B., Sweet, A.D., et al. (2022) Long-distance dispersal of pigeons and doves generated new ecological opportunities for host-switching and adaptive radiation by their parasites. Proceedings of the Royal Society B, 289, 20220042.

Broadinstitute/picard: A set of command line tools (in Java) for manipulating high-throughput sequencing (HTS) data and formats such as SAM/BAM/CRAM and VCF. [WWW Document]. (2022). URL https://github.com/broadinstitute/picard.

Buffalo, V. (2021) Quantifying the relationship between genetic diversity and population size suggests natural selection cannot explain Lewontin’s Paradox. eLife, 10, e67509.

Bush, A.O., Lafferty, K.D., Lotz, J.M. & Shostak, A.W. (1997) Parasitology Meets Ecology on Its Own Terms: Margolis et al. Revisited. The Journal of Parasitology, Parasitology Meets Ecology on Its Own Terms: Margolis et al. Revisited, 83.

Callaghan, C.T., Nakagawa, S. & Cornwell, W.K. (2021) Global abundance estimates for 9,700 bird species. Proceedings of the National Academy of Sciences, 118, e2023170118.

Capella-Gutiérrez, S., Silla-Martínez, J.M. & Gabaldón, T. (2009) trimAl: a tool for automated alignment trimming in large-scale phylogenetic analyses. Bioinformatics, 25, 1972–1973.

Carlson, C.J., Burgio, K.R., Dougherty, E.R., Phillips, A.J., Bueno, V.M., Clements, C.F., et al. (2017) Parasite biodiversity faces extinction and redistribution in a changing climate. Science Advances, 3, e1602422.

Carlson, C.J., Hopkins, S., Bell, K.C., Doña, J., Godfrey, S.S., Kwak, M.L., et al. (2020) A global parasite conservation plan. Biological Conservation, 250, 108596.

Charlesworth, B., Nordborg, M. & Charlesworth, D. (1997) The effects of local selection, balanced polymorphism and background selection on equilibrium patterns of genetic diversity in subdivided populations. Genetics Research, 70, 155–174.

Charlesworth, B., Charlesworth, D. & Barton, N.H. (2003) The effects of genetic and geographic structure on neutral variation. Annual Review of Ecology, Evolution, and Systematics, 34, 99–125.

Charlesworth, B. (2009) Effective population size and patterns of molecular evolution and variation. Nature Reviews Genetics, 10, 195–205.

Charlesworth, B. & Jensen, J.D. (2022) How Can We Resolve Lewontin’s Paradox? Genome Biology and Evolution, 14, evac096.

Chen, S., Zhou, Y., Chen, Y. & Gu, J. (2018) fastp: an ultra-fast all-in-one FASTQ preprocessor. Bioinformatics, 34, i884–i890.

Clayton, D.H. & Walther, B.A. (2001) Influence of host ecology and morphology on the diversity of Neotropical bird lice. Oikos, 94, 455–467.

Clayton, D.H. & Johnson, K.P. (2003) Linking coevolutionary history to ecological process: Doves and lice. Evolution, 57, 2335–2341.

Clayton, D.H., Bush, S. & Johnson, K.P. (2015) Coevolution of Life on Hosts: Integrating Ecology and History, Clayton, Bush, Johnson. University of Chicago Press.

Criscione, C.D. & Blouin, M.S. (2005) Effective sizes of macroparasite populations: a conceptual model. Trends in Parasitology, 21, 212–217.

Criscione, C.D., Poulin, R. & Blouin, M.S. (2005) Molecular ecology of parasites: elucidating ecological and microevolutionary processes. Molecular Ecology, 14, 2247–2257.

Danecek, P. & McCarthy, S.A. (2017) BCFtools/csq: haplotype-aware variant consequences. Bioinformatics, 33, btx100.

Danecek, P., Bonfield, J.K., Liddle, J., Marshall, J., Ohan, V., Pollard, M.O., et al. (2021) Twelve years of SAMtools and BCFtools. GigaScience, 10, giab008.

DiBlasi, E., Johnson, K.P., Stringham, S.A., Hansen, A.N., Beach, A.B., Clayton, D.H., et al. (2018) Phoretic dispersal influences parasite population genetic structure. Molecular Ecology, 27, 2770–2779.

Doña, J., Sweet, A.D. & Johnson, K.P. (2020) Comparing rates of introgression in parasitic feather lice with differing dispersal capabilities. Communications Biology, 3, 610.

Doña, J., Sweet, A.D., Johnson, K.P., Serrano, D., Mironov, S. & Jovani, R. (2017) Cophylogenetic analyses reveal extensive host-shift speciation in a highly specialized and host-specific symbiont system. Molecular Phylogenetics and Evolution, 115, 190–196.

Doña, J., Moreno-García, M., Criscione, C.D., Serrano, D. & Jovani, R. (2015) Species mtDNA genetic diversity explained by infrapopulation size in a host-symbiont system. Ecology and Evolution, 5, 5801–5809.

Doña, J. & Johnson, K.P. (2020) Assessing symbiont extinction risk using cophylogenetic data. Biological Conservation, 250, 108705.

Farrell, M.J., Park, A.W., Cressler, C.E., Dallas, T., Huang, S., Mideo, N., et al. (2021) The ghost of hosts past: impacts of host extinction on parasite specificity. Philosophical Transactions of the Royal Society B, 376, 20200351.

Felsenstein, J. (1985) Phylogenies and the Comparative Method. The American Naturalist, 125, 1–15.

Frankham, R. (1995) Effective population size/adult population size ratios in wildlife: a review. Genetical Research, 66, 95–107.

Freckleton, R.P., Harvey, P.H. & Pagel, M. (2002) Phylogenetic Analysis and Comparative Data: A Test and Review of Evidence. The American Naturalist, 160, 712–726.

H. Li (2022) seqtk Toolkit for processing sequences in FASTA/Q formats. URL https://github.com/lh3/seqtk

Haubold, B., Pfaffelhuber, P. & Lynch, M. (2010) mlRho –a program for estimating the population mutation and recombination rates from shotgun-sequenced diploid genomes. Molecular Ecology, 19, 277–284.B.

Harbison, C.W. & Clayton, D.H. (2011) Community interactions govern host-switching with implications for host–parasite coevolutionary history. Proceedings of the National Academy of Sciences, 108, 9525–9529.

Hedrick, P.W. & Garcia-Dorado, A. (2016) Understanding Inbreeding Depression, Purging, and Genetic Rescue. Trends in Ecology & Evolution, 31, 940–952.

Herrera, S.V., Johnson, K.P., Sweet, A.D., Ylinen, E., Kunnasranta, M. & Nyman, T. (2022) High levels of inbreeding with spatial and host-associated structure in lice of an endangered freshwater seal. Molecular Ecology, 31, 4593–4606.

Hoang, D.T., Chernomor, O., Haeseler, A. von, Minh, B.Q. & Vinh, L.S. (2017) UFBoot2: Improving the Ultrafast Bootstrap Approximation. Molecular Biology and Evolution, 35, 518–522.

Huyse, T., Poulin, R. & Théron, A. (2005) Speciation in parasites: a population genetics approach. Trends in Parasitology, 21, 469–475.

Johnson, K.P., Reed, D.L., Parker, S.L.H., Kim, D. & Clayton, D.H. (2007) Phylogenetic analysis of nuclear and mitochondrial genes supports species groups for Columbicola (Insecta: Phthiraptera). Molecular Phylogenetics and Evolution, 45, 506–518.

Johnson, K.P. & Weckstein, J.D. (2011) The Central American land bridge as an engine of diversification in New World doves. Journal of Biogeography, 38, 1069–1076.

Johnson, K.P., Dietrich, C.H., Friedrich, F., Beutel, R.G., Wipfler, B., Peters, R.S., et al. (2018) Phylogenomics and the evolution of hemipteroid insects. Proceedings of the National Academy of Sciences, 115, 201815820.

Johnson, K.P., Weckstein, J.D., Herrera, S.V. & Doña, J. (2021) The interplay between host biogeography and phylogeny in structuring diversification of the feather louse genus Penenirmus. Molecular Phylogenetics and Evolution, 107297.

Johnson, K.P., Matthee, C. & Doña, J. (2022) Phylogenomics reveals the origin of mammal lice out of Afrotheria. Nature Ecology & Evolution, 6, 1205–1210.

Katoh, K. & Standley, D.M. (2013) MAFFT Multiple Sequence Alignment Software Version 7: Improvements in Performance and Usability. Molecular Biology and Evolution, 30, 772–780.

Kimura, M. (1971) Theoretical foundation of population genetics at the molecular level. Theoretical Population Biology, 2, 174–208.

Kirkness, E.F., Haas, B.J., Sun, W., Braig, H.R., Perotti, M.A., Clark, J.M., et al. (2010) Genome sequences of the human body louse and its primary endosymbiont provide insights into the permanent parasitic lifestyle. Proceedings of the National Academy of Sciences, 107, 12168–12173.

Larsen, L. -K., Pélabon, C., Bolstad, G.H., Viken, Å., Fleming, I.A. & Rosenqvist, G. (2011) Temporal change in inbreeding depression in life-history traits in captive populations of guppy (Poecilia reticulata): evidence for purging? Journal of Evolutionary Biology, 24, 823–834.

Lewontin, R. C. (1974) The genetic basis of evolutionary change (Vol. 560). Columbia University Press, New York.

Li, H. & Durbin, R. (2009) Fast and accurate short read alignment with Burrows–Wheeler transform. Bioinformatics, 25, 1754–1760.

Lynch, M. (2008) Estimation of Nucleotide Diversity, Disequilibrium Coefficients, and Mutation Rates from High-Coverage Genome-Sequencing Projects. Molecular Biology and Evolution, 25, 2409–2419.

Malenke, J.R., Johnson, K.P. & Clayton, D.H. (2009) Host Specialization Differentiates Cryptic Species of Feather-Feeding Lice. Evolution, 63, 1427–1438.

Martins, E.P. & Hansen, T.F. (1997) Phylogenies and the Comparative Method: A General Approach to Incorporating Phylogenetic Information into the Analysis of Interspecific Data. The American Naturalist, 149, 646–667.

Martinů, J., Hypša, V. & Štefka, J. (2018) Host specificity driving genetic structure and diversity in ectoparasite populations: Coevolutionary patterns in Apodemus mice and their lice. Ecology and Evolution, 8, 10008–10022.

Matthews, A.E., Klimov, P.B., Proctor, H.C., Dowling, A.P.G., Diener, L., Hager, S.B., et al. (2018) Cophylogenetic assessment of New World warblers (Parulidae) and their symbiotic feather mites (Proctophyllodidae). Journal of Avian Biology, 49, jav–01580.

Meyer, M., Kircher, M., Gansauge, M.-T., Li, H., Racimo, F., Mallick, S., et al. (2012) A High-Coverage Genome Sequence from an Archaic Denisovan Individual. Science, 338, 222–226.

Minh, B.Q., Schmidt, H.A., Chernomor, O., Schrempf, D., Woodhams, M.D., Haeseler, A. von, et al. (2020) IQ-TREE 2: New models and efficient methods for phylogenetic inference in the genomic era. Molecular Biology and Evolution, 37, 1530–1534.

Moya, R.S. de, Allen, J.M., Sweet, A.D., Walden, K.K.O., Palma, R.L., Smith, V.S., et al. (2019) Extensive host-switching of avian feather lice following the Cretaceous-Paleogene mass extinction event. Communications Biology, 2, 445.

Nadler, S.A., Hafner, M.S., Hafner, J.C. & Hafner, D.J. (1990) Genetic differentiation among chewing louse populations (Mallophaga: Trichodectidae) in a pocket gopher contact zone (Rodentia: Geomyidae). Evolution, 44, 942–951.

Nee, S., Read, A.F., Greenwood, J.J.D. & Harvey, P.H. (1991) The relationship between abundance and body size in British birds. Nature, 351, 312–313.

Nieminen, M., Rita, H. & Uuvana, P. (1999) Body size and migration rate in moths. Ecography, 22, 697–707.

Nowak, J., Sweet, A., Weckstein, J. & Johnson, K. (2019) A molecular phylogenetic analysis of the genera of fruit doves and allies using dense taxonomic sampling. Illinois Natural History Survey Bulletin, 42, 2019001.

Pagel, M. (1999) Inferring the historical patterns of biological evolution. Nature, 401, 877–884.

Paradis, E. & Schliep, K. (2018) ape 5.0: an environment for modern phylogenetics and evolutionary analyses in R. Bioinformatics, 35, 526–528.

Pinheiro, J., Bates, D., DebRoy, S., Sarkar, D. & Team, R.C. (2020) nlme: Linear and Nonlinear Mixed Effects Models.

Poulin, R. (1999) Body size vs abundance among parasite species: positive relationships? Ecography, 22, 246–250.

Poulin, R. (2007) Evolutionary Ecology of Parasites. Princeton University Press, Princeton, New Jersey.

Revell, L.J. (2012) phytools: an R package for phylogenetic comparative biology (and other things). Methods in Ecology and Evolution, 3, 217–223.

Robinson, J.A., Brown, C., Kim, B.Y., Lohmueller, K.E. & Wayne, R.K. (2018) Purging of Strongly Deleterious Mutations Explains Long-Term Persistence and Absence of Inbreeding Depression in Island Foxes. Current Biology, 28, 3487–3494.e4.

Rozsa, L. (1997) Patterns in the Abundance of Avian Lice (Phthiraptera: Amblycera, Ischnocera). Journal of Avian Biology, 28, 249.

Santichaivekin, S., Yang, Q., Liu, J., Mawhorter, R., Jiang, J., Wesley, T., et al. (2020) eMPRess: a systematic cophylogeny reconciliation tool. Bioinformatics.

Slater, G.S.C. & Birney, E. (2005) Automated generation of heuristics for biological sequence comparison. BMC Bioinformatics, 6, 31–31.

Strobel, H.M., Hays, S.J., Moody, K.N., Blum, M.J. & Heins, D.C. (2019) Estimating effective population size for a cestode parasite infecting three-spined sticklebacks. Parasitology, 146, 883–896.

Sweet, A.D., Maddox, J.D. & Johnson, K.P. (2017) A complete molecular phylogeny of Claravis confirms its paraphyly within small New World ground-doves (Aves: Peristerinae) and implies multiple plumage state transitions. Journal of Avian Biology, 48, 459–464.

Sweet, A.D. & Johnson, K.P. (2018) The role of parasite dispersal in shaping a host–parasite system at multiple evolutionary scales. Molecular Ecology, 27, 5104–5119.

To, T.-H., Jung, M., Lycett, S. & Gascuel, O. (2016) Fast Dating Using Least-Squares Criteria and Algorithms. Systematic Biology, 65, 82–97.

Valk, T. van der, Manuel, M. de, Marques-Bonet, T. & Guschanski, K. (2021) Estimates of genetic load suggest frequent purging of deleterious alleles in small populations. bioRxiv, 696831.

Waples, R.S. (2002) Definition and estimation of effective population size in the conservation of endangered species. In Population Viability Analysis (ed. by Beissinger, S.R. & McCullough, D.R.). University of Chicago Press, Chicago.

White, E.P., Ernest, S.K.M., Kerkhoff, A.J. & Enquist, B.J. (2007) Relationships between body size and abundance in ecology. Trends in Ecology & Evolution, 22, 323–330.

Wright, S. (1943) Isolation by distance. Genetics, 28, 114–138.

Zhang, C., Rabiee, M., Sayyari, E. & Mirarab, S. (2018) ASTRAL-III: polynomial time species tree reconstruction from partially resolved gene trees. BMC Bioinformatics, 19, 15

