## Supplementary material for "Host body size, not host population size, predicts genome-wide effective population size of parasites": Figure S1

**Fig S1. Cladogram of *Columbicola* host species used for the cophylogenetic analyses. The tree file can be found in (reserved DOI: 10.6084/m9.figshare.21269640; private link for review: https://figshare.com/s/2f2de5dc909155da815a)**

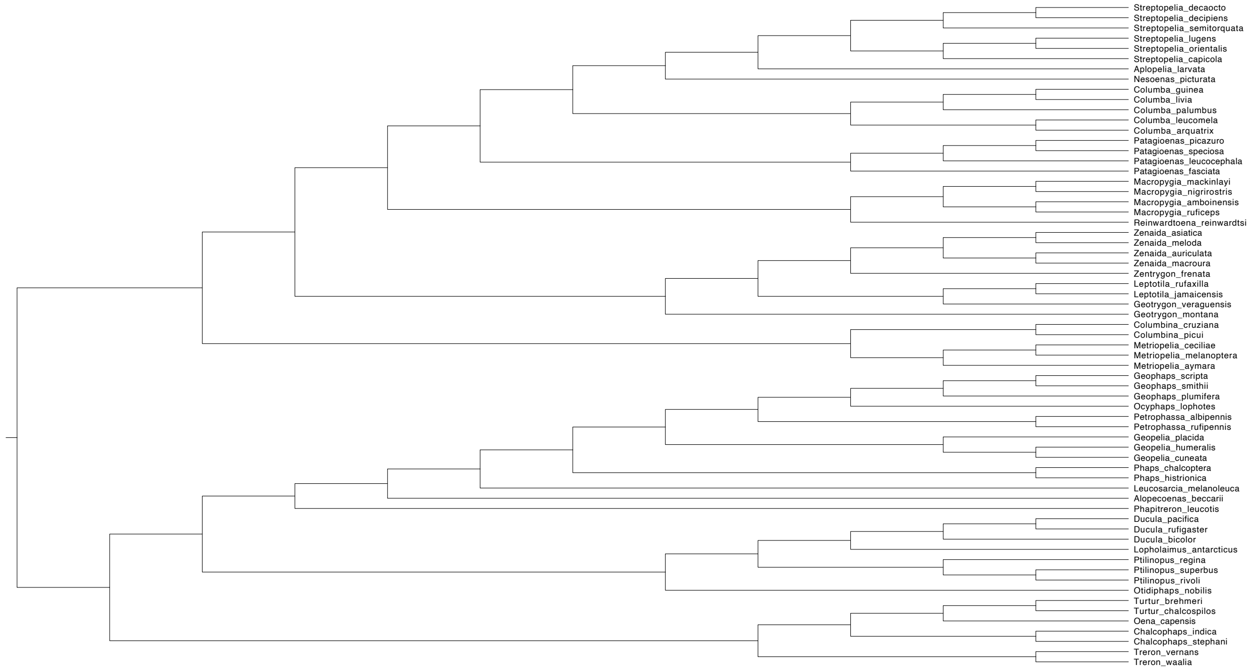
