## Supplementary material for "Host body size, not host population size, predicts genome-wide effective population size of parasites": Figure S2

**Fig S2. Time-calibrated tree of *Columbicola* used for the phylogenetic comparative analyses. Numbers on branches are ultrafast bootstrap values from IQ-TREE. The full tree can be found in (reserved DOI: 10.6084/m9.figshare.21269640; private link for review: https://figshare.com/s/2f2de5dc909155da815a)**

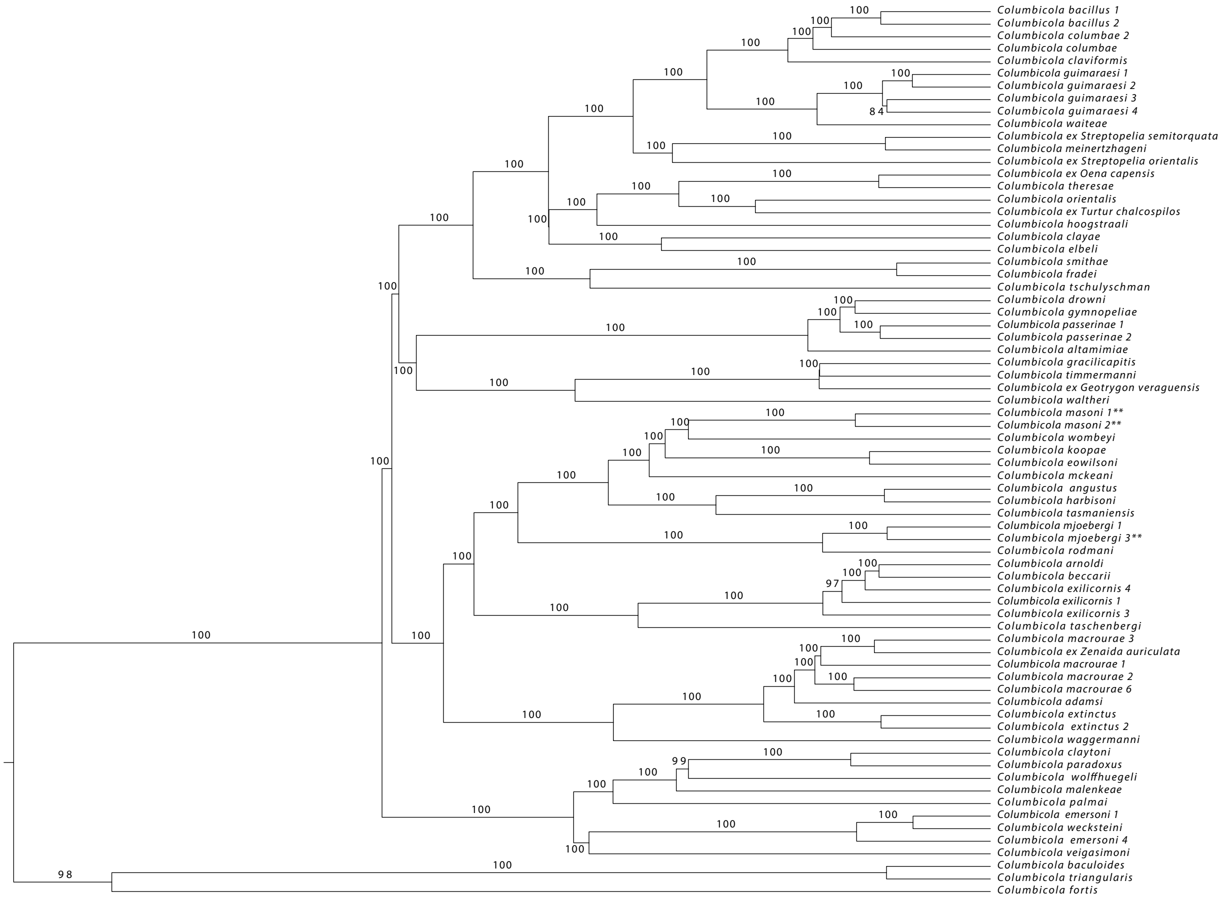
