## Supplementary material for "Host body size, not host population size, predicts genome-wide effective population size of parasites": Figure S3

### Fig S3. Scatter plots showing the relationship between theta and host population size (both untransformed and log transformed).

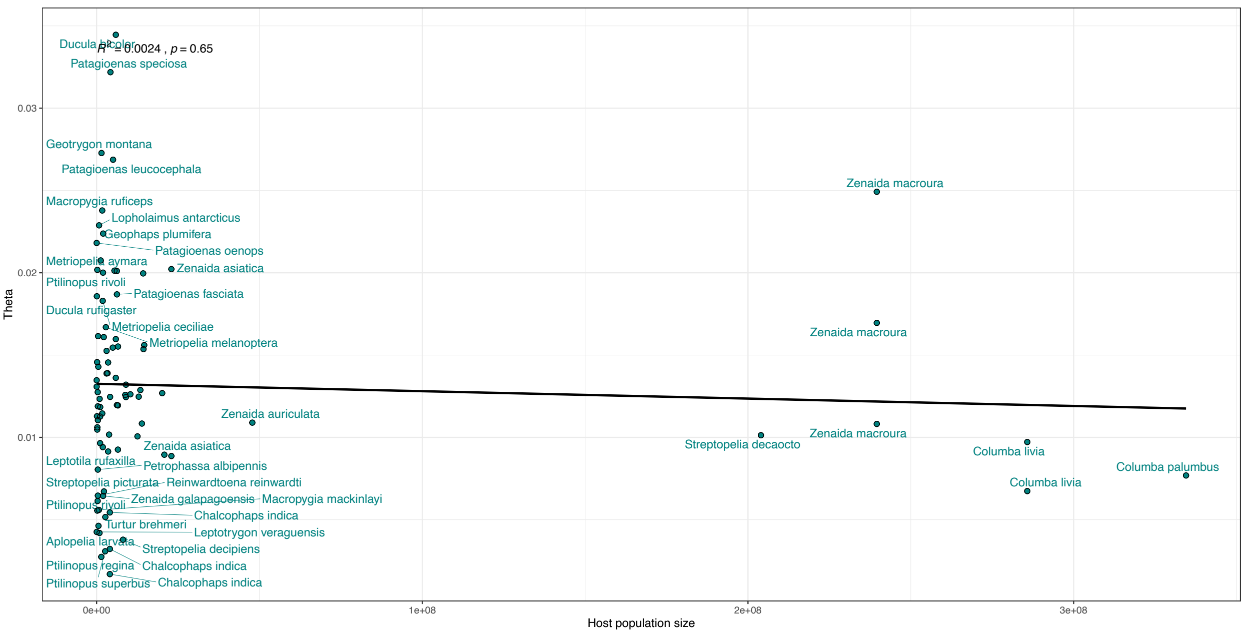


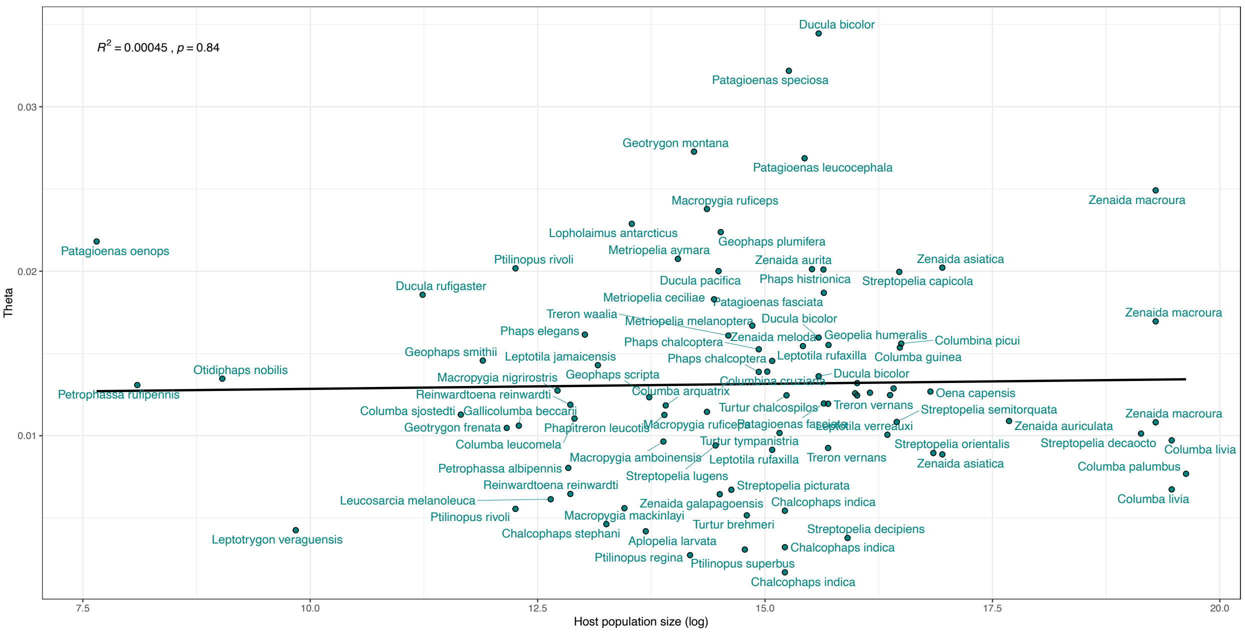
