## Supplementary material for "Host body size, not host population size, predicts genome-wide effective population size of parasites": Table S2

Table S2. Cospeciation events used as calibration points in study.

| **Louse species (1)** | **Host species (1)** | **Louse species (2)** | **Host species (2)** | **Minimum age (95 % CI, mya)** | **Maximum age (95 % CI, mya)** |
| --- | --- | --- | --- | --- | --- |
| *Columbicola bacillus* | *Streptopelia decaocto* | *Columbicola bacillus* | *Streptopelia decipiens* | 1 | 13 |
| *Columbicola macrourae* | *Zenaida meloda* | *Columbicola macrourae* | *Zenaida asiatica* | 1 | 10 |
| *Columbicola passerinae* | *Columbina picui* | *Columbicola passerinae* | *Columbina cruziana* | 1 | 24 |
| *Columbicola drowni* | *Metropelia melanoptera* | *Columbicola gymnopeliae* | *Metriopelia ceciliae* | 2 | 29 |
| *Columbicola koopae* | *Geophaps scripta* | *Columbicola eowilsoni* | *Geophaps smithii* | 1 | 10 |
| *Columbicola masoni* | *Petrophassa rufipennis* | *Columbicola masoni* | *Petrophassa albipennis* | 2 | 17 |
| *Columbicola clayae* | *Treron waalia* | *Columbicola elbeli* | *Treron vernans* | 2 | 35 |
