## Supplementary material for "Host body size, not host population size, predicts genome-wide effective population size of parasites": Table S1

**Table S1. Samples in study**

| **Louse species** | **Bird host** | **Country** | **# Reads (millions after trimming)** | **# Loci** | **# SNPs retained after filtering** | **NCBI SRA accession** |
| --- | --- | --- | --- | --- | --- | --- |
| *Columbicola paradoxus* | *Lopholaimus antarcticus* | Australia | 38.4 | 2303 | 128943 | SRR3161944 |
| *Columbicola fortis* | *Otidiphaps nobilis* | Papua New Guinea | 87.4 | 2349 | 156295 | SRR3161925 |
| *Columbicola claytoni* | *Ducula rufigaster* | Papua New Guinea | 70.8 | 2347 | 176689 | SRR3161916 |
| *Columbicola claviformis* | *Columba palumbus* | United Kingdom | 57 | 2351 | 177699 | SRR3161920 |
| *Columbicola wolffhuegeli* | *Ducula bicolor* | Australia | 148.9 | 2342 | 128637 | SRR8145890 |
| *Columbicola waiteae* | *Columba leucomela* | Australia | 65.4 | 2340 | 175723 | SRR3161940 |
| *Columbicola palmai* | *Leucosarcia melanoleuca* | Australia | 17.2 | 2336 | 173282 | SRR16503055 |
| *Columbicola triangularis* | *Patagioenas picazuro* | Argentina | 105.2 | 2328 | 136312 | SRR3161964 |
| *Columbicola malenkeae* | *Ducula pacifica* | Vanuatu | 80.7 | 2341 | 175159 | SRR3161956 |
| *Columbicola columbae* | *Columba livia* | United States of America | 74 | 2337 | 177658 | SRR8177102 |
| *Columbicola extinctus 2* | *Patagioenas fasciata* | Peru | 169.9 | 2344 | 90992 | SRR8145888 |
| *Columbicola extinctus* | *Patagioenas fasciata* | United States of America | 46.7 | 2346 | 175236 | SRR3161924 |
| *Columbicola meinertzhageni* | *Columba arquatrix* | Malawi | 118.6 | 2323 | 170125 | SRR16503057 |
| *Columbicola tschulyschman* | *Columba livia* | United States of America | 43.5 | 2344 | 172843 | SRR3161959 |
| *Columbicola angustus* | *Phaps chalcoptera* | Australia | 78.1 | 2336 | 174679 | SRR8145885 |
| *Columbicola tasmaniensis* | *Phaps chalcoptera* | Australia | 67.2 | 2343 | 175771 | SRR3161947 |
| *Columbicola waltheri* | *Geotrygon frenata* | Peru | 58.7 | 2327 | 173979 | SRR3161933 |
| *Columbicola adamsi* | *Patagioenas speciosa* | Mexico | 41.5 | 2330 | 165065 | SRR3161912 |
| *Columbicola columbae 2* | *Columba guinea* | South Africa | 37.9 | 2341 | 177143 | SRR3161917 |
| *Columbicola harbisoni* | *Phaps histrionica* | Australia | 70.2 | 2329 | 171283 | SRR3161949 |
| *Columbicola clayae* | *Treron waalia* | Ghana | 40.8 | 2334 | 150546 | SRR3161934 |
| *Columbicola taschenbergi* | *Reinwardtoena reinwardti* | Papua New Guinea | 50.2 | 2346 | 164171 | SRR16503052 |
| *Columbicola waggermanni* | *Patagioenas leucocephala* | Jamaica | 121.3 | 2319 | 141157 | SRR16503050 |
| *Columbicola sp.* | *Streptopelia semitorquata* | Ghana | 50.8 | 2323 | 172990 | SRR3161935 |
| *Columbicola sp.* | *Streptopelia orientalis* | China | 61.8 | 2331 | 172534 | SRR3161951 |
| *Columbicola macrourae 6* | *Zenaida meloda* | Peru | 86.9 | 2347 | 175948 | SRR3161971 |
| *Columbicola koopae* | *Geophaps scripta* | Australia | 84.6 | 2345 | 170503 | SRR3161958 |
| *Columbicola mckeani* | *Ocyphaps lophotes* | Australia | 53 | 2261 | 114534 | SRR3161929 |
| *Columbicola eowilsoni* | *Geophaps smithii* | Australia | 71.3 | 2343 | 173044 | SRR3161960 |
| *Columbicola bacillus 2* | *Streptopelia decipiens* | Uganda | 85.6 | 2349 | 177546 | SRR3161967 |
| *Columbicola hoogstraali* | *Streptopelia picturata* | Madagascar | 63.9 | 2343 | 175954 | SRR3161968 |
| *Columbicola orientalis* | *Streptopelia lugens* | Malawi | 82.4 | 2340 | 176721 | SRR3161972 |
| *Columbicola bacillus 1* | *Streptopelia decaocto* | Netherlands | 69.2 | 2342 | 169738 | SRR3161950 |
| *Columbicola masoni 2*** | *Petrophassa rufipennis* | Australia | 43.8 | 2334 | 171177 | SRR3161946 |
| *Columbicola sp.* | *Leptotrygon veraguensis* | Panama | 118.7 | 2340 | 175914 | SRR16503054 |
| *Columbicola fradei* | *Aplopelia larvata* | Malawi | 53.9 | 2343 | 176300 | SRR3161954 |
| *Columbicola gracilicapitis* | *Leptotila jamaicensis* | Mexico | 73.9 | 2336 | 174728 | SRR3161913 |
| *Columbicola theresae* | *Streptopelia capicola* | South Africa | 61.6 | 2340 | 176768 | SRR3161915 |
| *Columbicola macrourae 2* | *Zenaida asiatica* | United States of America | 73.7 | 2339 | 175505 | SRR3161952 |
| *Columbicola timmermanni* | *Leptotila rufaxilla* | Guyana | 122.5 | 2328 | 175374 | SRR16503051 |
| *Columbicola wecksteini* | *Ptilinopus rivoli* | Papua New Guinea | 131.2 | 2339 | 164396 | SRR16503027 |
| *Columbicola exilicornis 1* | *Macropygia amboinensis* | Papua New Guinea | 82.9 | 2346 | 175511 | SRR3161939 |
| *Columbicola macrourae 1* | *Geotrygon montana* | Mexico | 51.3 | 2341 | 170670 | SRR3161914 |
| *Columbicola elbeli* | *Treron vernans* | Malaysia | 118.1 | 2328 | 162380 |  |
| *Columbicola rodmani* | *Geopelia humeralis* | Australia | 67.4 | 2333 | 176613 | SRR3161918 |
| *Columbicola masoni 1*** | *Petrophassa albipennis* | Australia | 88 | 2349 | 152169 | SRR3161937 |
| *Columbicola baculoides* | *Zenaida macroura* | United States of America | 102.9 | 2339 | 153037 | SRR13159337 |
| *Columbicola macrourae 3* | *Zenaida macroura* | United States of America | 64.1 | 2330 | 175767 | SRR3161953 |
| *Columbicola guimaraesi 1* | *Chalcophaps indica* | Vanuatu | 42 | 2342 | 175584 | SRR3161955 |
| *Columbicola guimaraesi 2* | *Chalcophaps indica* | Australia | 145.8 | 2341 | 93343 | SRR16503017 |
| *Columbicola guimaraesi 4* | *Chalcophaps indica* | China | 121.1 | 2337 | 177542 | SRR16503016 |
| *Columbicola smithae* | *Turtur brehmeri* | Ghana | 49.4 | 2341 | 175118 | SRR3161936 |
| *Columbicola guimaraesi 3* | *Chalcophaps stephani* | Papua New Guinea | 60.6 | 2349 | 177494 | SRR3161927 |
| *Columbicola veigasimoni* | *Phapitreron leucotis* | Philippines | 77.4 | 2338 | 174478 | SRR3161919 |
| *Columbicola emersoni 1* | *Ptilinopus superbus* | Australia | 87 | 2333 | 175568 | SRR8145886 |
| *Columbicola emersoni 4* | *Ptilinopus regina* | Australia | 90.1 | 2334 | 175478 | SRR8145887 |
| *Columbicola sp.* | *Zenaida auriculata* | Ecuador | 115.3 | 2342 | 176011 | SRR16503053 |
| *Columbicola drowni* | *Metriopelia melanoptera* | Argentina | 55.7 | 2344 | 176801 | SRR3161922 |
| *Columbicola wombeyi* | *Geophaps plumifera* | Australia | 40.6 | 2333 | 168808 | SRR3161943 |
| *Columbicola exilicornis 4* | *Macropygia mackinlayi* | Vanuatu | 76 | 2350 | 176796 | SRR3161945 |
| *Columbicola arnoldi* | *Macropygia nigrirostris* | Papua New Guinea | 86.5 | 2344 | 176274 | SRR3161961 |
| *Columbicola exilicornis 3* | *Macropygia ruficeps* | Malaysia | 33 | 2325 | 151023 | SRR3161962 |
| *Columbicola beccarii* | *Gallicolumba beccarii* | Solomon Islands | 50.6 | 2349 | 171261 | SRR3161941 |
| *Columbicola altamimiae* | *Metriopelia aymara* | Argentina | 48.9 | 2342 | 174347 | SRR3161921 |
| *Columbicola gymnopeliae* | *Metriopelia ceciliae* | Peru | 38 | 2340 | 166272 | SRR3161923 |
| *Columbicola sp.* | *Turtur chalcospilos* | Malawi | 67.2 | 2337 | 171128 | SRR3161970 |
| *Columbicola mjoebergi 3*** | *Geopelia placida* | Australia | 83.7 | 2346 | 175726 | SRR3161942 |
| *Columbicola passerinae 1* | *Columbina picui* | Argentina | 45.4 | 2326 | 151243 | SRR3161931 |
| *Columbicola passerinae 2* | *Columbina cruziana* | Peru | 54.4 | 2348 | 174325 | SRR3161930 |
| *Columbicola sp.* | *Oena capensis* | Madagascar | 112.2 | 2329 | 174275 | SRR8177104 |
| *Columbicola mjoebergi 1* | *Geopelia cuneata* | Australia | 101.6 | 2347 | 145368 | SRR3161957 |
